# MUTYH activity maintains telomere stability in response to chronic telomeric 8-oxoguanine damage in cancer cells

**DOI:** 10.64898/2026.07.31.742130

**Authors:** Mariarosaria De Rosa, Theresa Heidenreich, Libby Childs, Benura Azeroglu, Sneh M Toprani, Nader Aryamanesh, Pablo Galaviz, Hilda A Pickett, Eros Lazzerini Denchi, Zachary Nagel, Patricia L. Opresko

## Abstract

Telomeres are highly susceptible to oxidative DNA damage, particularly 8-oxoguanine (8-oxoG), which is processed by glycosylase-initiated base excision repair (BER). OGG1 removes 8-oxoG opposite C, and MUTYH removes A misinserted opposite 8-oxoG to prevent mutations. While OGG1 has an established role in telomere protection, the contribution of MUTYH to telomere stability in cancer cells after oxidative DNA damage remains poorly understood. Using a chemoptogenetic system to induce targeted 8-oxoG lesions specifically at telomeres in HeLa cancer cells, we demonstrate that MUTYH is required to prevent telomere shortening, telomere loss, and genomic instability after chronic damage. Yet, telomere damage in MUTYH-deficient cells does not cause sustained DNA damage signaling or reduced cellular proliferation. Whole-genome sequencing further reveals enrichment of G to T transversions within telomeric repeats in MUTYH-deficient cells, consistent with increased mutagenesis due to unrepaired 8-oxoG:A mispairs. Combined loss of MUTYH and OGG1 rescues damage-induced telomere aberrations and genomic instability, implicating BER-generated single-strand break (SSB) intermediates as major contributors to telomere instability. In agreement, exo-FISH and S1-END-seq analyses reveal that repair-proficient cells rapidly accumulate SSB intermediates after damage, which are later resolved, whereas glycosylase-deficient cells exhibit SSBs at later time points. Together, these findings identify MUTYH as a critical guardian of telomere integrity during chronic oxidative stress and provide insight into how defective BER at telomeres contributes to genomic instability in cancer cells, with implications for cancers associated with MUTYH deficiency and mutations.

## INTRODUCTION

Telomeres are specialized nucleoprotein structures that protect chromosome ends from being recognized as DNA double-strand breaks (DSBs) and are essential for preserving genome stability [1]. Their repetitive, G-rich DNA sequence renders telomeres particularly susceptible to oxidative damage, especially the formation of 8-oxoguanine (8-oxoG), which is among the most prevalent base lesions. 8-oxoG arises frequently under conditions of oxidative stress, a major driver of telomere dysfunction and shortening [2]. Accumulation of oxidative DNA damage at telomeres interferes with normal replication, compromises shelterin binding, and promotes telomere fragility and loss, chromosome end-to-end fusions, and genome instability, which are hallmarks of both cellular senescence and cancer progression [3–5].

Base excision repair (BER) represents the primary pathway responsible for repairing oxidative DNA damage, including 8-oxoG [6]. Within this pathway, the DNA glycosylases 8-oxoguanine glycosylase 1 (OGG1) and MutY homolog (MUTYH) act in a coordinated manner to prevent mutagenesis: OGG1 excises 8-oxoG when paired with cytosine, whereas MUTYH removes adenines misincorporated opposite 8-oxoG during DNA replication. Previous work from our laboratory and others has demonstrated that OGG1 plays a critical role in preserving telomere integrity following oxidative damage in cancer cells [4,7]. Using a chemoptogenetic approach to induce targeted 8-oxoG lesions specifically at telomeres, we showed that failure to repair chronic telomeric 8-oxoG damage promotes telomere shortening, loss and crisis [4]. More recently, we reported that OGG1 and MUTYH activity contribute to cellular senescence induction in non-diseased human cells in response to telomeric 8-oxoG, highlighting that the biological consequences of oxidative base damage and its repair at telomeres are context-dependent [8].

MUTYH is a critical genome maintenance protein with particular relevance to cancer biology. Germline biallelic mutations in MUTYH cause MUTYH-associated polyposis (MAP), a hereditary cancer predisposition syndrome characterized by a markedly increased lifetime risk of colorectal cancer [9]. Notably, the most prevalent pathogenic MUTYH mutations impair either its glycosylase activity or its ability to bind 8-oxoG. This is a crucial initial step for the removal of the misinserted adenines to prevent G:C to T:A transversion mutations from being permanently fixed into the genome [10–12]. Beyond inherited disease, somatic MUTYH mutations are detected in a wide spectrum of sporadic tumors, underscoring its relevance in tumorigenesis [13,14]. Mouse models further support a unique tumor-suppressive role for MUTYH. Although loss of either OGG1 or MUTYH increases oxidative DNA damage and mutagenesis, only Mutyh-deficient mice develop tumors following chronic oxidative stress, suggesting that MUTYH performs functions that extend beyond only mutation avoidance [15]. Despite the strong association between MUTYH deficiency and cancer, its role in maintaining telomere stability in cancer cells remains poorly defined. A few studies have reported MUTYH recruitment to telomeres following oxidative stress in mouse embryonic fibroblasts (MEFs) or HEK-293T cells [16], and interactions between MUTYH and telomere-associated factors have been proposed [17]. MUTYH-deficient HEK-293T cells show extra-chromosomal telomeres and increased telomere fusions after H_2_O_2_ treatment [17]. However, the consequences of MUTYH loss on telomere stability after oxidative damage in cancer cells have been largely unexplored.

Here, we addressed this gap in knowledge by targeting 8-oxoG lesions at telomeres in cancer cells rendered deficient for MUTYH. This approach allowed us to dissect the specific contribution of MUTYH to telomere maintenance under conditions of chronic telomeric oxidative stress, independent of pleiotropic effects caused by global oxidative stress. We observed that MUTYH loss leads to progressive telomere shortening, increased telomere loss, and elevated genomic instability in HeLa long telomere (LT) cells chronically exposed to telomeric 8-oxoG. Notably, these defects occur without a corresponding reduction in cellular proliferation, suggesting that MUTYH deficiency enhances the ability of cancer cells to tolerate increased chromosomal instability driven by oxidative telomeric damage. Furthermore, combined deficiency of MUTYH and OGG1 suppresses telomere aberrations induced by chronic oxidative damage, implicating BER intermediates as key drivers of telomere instability. Together, our findings uncover a previously unappreciated role for MUTYH in safeguarding telomere stability in cancer cells under chronic oxidative stress and provide mechanistic insight into how inefficient BER completion at telomeres may contribute to genome instability and cancer progression.

## MATERIALS AND METHODS

### Cell culture and cell lines generation

HeLa FAP-mCER-TRF1 (HeLa FAP-TRF1) wild type cells were previously described [4] and were cultured at 5% oxygen in DMEM supplemented with 10% FBS, 1% penicillin/streptomycin and 500 µg/ml G418 (Gibco). Knock out (KO) cell lines were obtained by transfecting HEK293T cells (ATCC) with pLentiCRISPR V2 vectors (GenScript) expressing S. pyogenes Cas9, guide RNAs designed and validated for uniquely targeting the human MUTYH gene [18], and with Mission Packaging Mix (Sigma) to produce lentivirus. The lentivirus collected 48h and 72h post transfection was used to infect the recipient cells, and then selected as described below. MUTYH KO cells were obtained by infection of HeLa FAP-TRF1WT with lentivirus expressing respectively guide RNAs (gRNAs) targeting MUTYH intron 2 (gRNA1: CTTGGTCGTACCAGCTTAGC for clone 1.5; gRNA2: ACTGTGATCAACTACTATAC for clone 2.22), and selected with 1.5 µg/ml Puromycin (Gibco). OGG1 KO/MUTYH KO (DKO) cells were obtained by infection of HeLa FAP-TRF1 OGG1 KO c3.14 [4] with lentivirus expressing a gRNA targeting MUTYH exon 2 (gRNA5: GCATGCTAAGAACAACAGTC for clones 5.1 and 5.4), and were selected with 10 µg/ml Blasticidin S HCl (Gibco). After selection and death of uninfected cells, the infected cells were single cell cloned. Individual clones were selected based on abolished expression of MUTYH, which was determined by western blotting using MUTYH antibody (Abnova #H00004595-M01).

### Cell treatments

Telomere localized singlet oxygen was generated by treating cells overexpressing FAP-TRF1 with malachite green di-iodinated (MG2I) dye and light, as described previously [4]. Briefly, cells were seeded and incubated overnight. The next day cells were incubated in OptiMEM media (Gibco) at 37 ° C for 15 min before adding 100 nM MG2I dye for another 15 min. Cells were then exposed in the lightbox to a high-intensity 660 nm LED light at 100 mW/cm^2^ for 5 min (unless indicated otherwise) to trigger excitation of the FAP-bound MG2I dye and the production of singlet oxygen. For chronic exposure to telomeric singlet oxygen, 2×10^5^ HeLa FAP-TRF1 cells were plated and treated with 5 min dye and light as described above, for three consecutive days and harvested/reseeded every fourth day for a total of 24 days and 18 exposures.

### Western blotting

Cells were washed twice with PBS and then lysed directly in the dish, on ice with RIPA buffer (Santa Cruz) supplemented with PMSF (1 nM), 1x Roche Protease and Phosphatase Inhibitors, and Benzonase (Sigma E8263; 1:500) for 15 min and then incubated at 37 °C for 10 min, before spinning down at 15,000 rpm for 10 minutes at 4 °C. Protein concentrations were determined with the BCA assay (Pierce) and 20-30 μg of protein was electrophoresed on 4–12% Bis-Tris gels (Thermo) before semi-dry transfer to nitrocellulose membranes (GE Healthcare). Red Ponceau staining was performed to ensure even transfer of proteins onto membranes, which were then washed in TBS-T and blocked in 5% milk, and blotted with primary and secondary HRP antibodies. The signal was detected by ECL detection and acquired on the iBright^TM^ FL1500 Imaging System (Thermo Fisher Scientific). Antibodies used were anti-MUTYH mouse monoclonal (Abnova #H00004595-M01, WB dilution 1:1000), anti-OGG1 rabbit monoclonal (Cell Signaling #46271S, WB dilution 1:500), anti-alpha-tubulin mouse monoclonal (Millipore #05-829, WB dilution 1:5000).

### Measurement of DNA Repair Capacity Using Fluorescence Multiplex based Host Cell Reactivation (FM-HCR) assays

Reporter plasmids were prepared, and the Fluorescence Multiplex based Host Cell Reactivation (FM-HCR) assays were performed as previously described [19]. All FM-HCR reporter constructs were based on the pMax vector backbone. Plasmid reporters containing site-specific 8-oxoG-C lesions and A-8-oxoG lesions were generated in pMax_mOrange and pMax_mPlum vectors, respectively, as described previously [19]. Two reporter plasmid cocktails were prepared for FM-HCR analyses. The damaged reporter cocktail consisted of 100 ng each of pMax_mOrange_8-oxoG-C, pMax_mPlum_A-8-oxoG along with pMax_GFP as transfection efficiency control. The undamaged reporter cocktail contained 100 ng each of pMax_GFP, pMax_mOrange and pMax_mPlum. For transfection experiments, 2×10^5^ cells were seeded per well in 12-well plates and allowed to adhere overnight. On the day of transfection, the culture medium was replaced with 1 mL of fresh complete DMEM culture medium, and cells were transfected with 1.5 μg of either the undamaged or damaged plasmid cocktails using Lipofectamine 3000 (Thermo Fisher Scientific), following the manufacturer’s instructions. Untransfected cells were included as negative controls, as previously described [20]. After 24h incubation at 37°C in a humidified atmosphere containing 5% CO_2_, cells were trypsinized, resuspended in complete DMEM culture medium and analyzed using an Attune NxT flow cytometer (Thermo Fisher Scientific). Compensation and gating were established using untransfected cells and single-color controls. Reporter expression was normalized to transfection efficiency and pathway-specific DNA repair capacity was calculated as previously described [19,21].

### Population doubling measurement

The population doubling (PD) values were calculated using the mathematical formula PD = [(ln(N2)) - (ln(N1))] / ln(2), where N1 is the initial number of cells plated and N2 the final number of cells counted. PD curves were obtained using the sum of the individual PDs calculated every 4 days.

### Telomere fluorescence *in situ* hybridization (FISH) on metaphase spreads

Chromosome spreads were prepared by incubating cells with 0.05 μg/ml colcemid for 2 h prior to harvesting with trypsin. Cells were incubated with 75 mM KCl for 13 min at 37 °C and fixed in methanol and glacial acetic acid (3:1). Cells were dropped on washed slides and dried overnight before fixation in 4% formaldehyde. Slides were treated with RNaseA and Pepsin at 37 °C, and then dehydrated. FISH was performed by diluting the telomeric PNA probe 1:100 (PNA Bio, F1004, TelC-Alexa 488, 3xCCCTAA) and a Pan-centromere probe 1:200 (PNA Bio, F3005, CENPB-Cy5, ATTCGTTGGAAACGGGA) in hybridization buffer (70% formamide, 10 mM Tris HCl pH 7.5, 1x Maleic Acid buffer, 1x MgCl2 buffer) which was then boiled for 5 min at 85 °C before returning to ice. Coverslips were incubated with the complete hybridization buffer for 10 min on a hot plate at 75 °C and then kept at 4 °C overnight in dark humid chambers. The following day, after two washes in hybridization wash buffer (70% formamide, 10 mM tris HCl pH 7.5), slides were rinsed in water before dehydrating in sequential ethanol washes (70%, 90% and 100%). Once completely dry, slides were stained with DAPI-containing mounting with ProLong™ Diamond Antifade Mountant (Thermo Fisher). The Nikon NIS Elements AR software was used for qFISH analyses to measure single telomere signal intensities. Each metaphase was isolated as single ROI and telomeres were selected as objects creating a binary layer based on their intensities. The binary object intensities were then collected and separately binned into intervals of 2500 to obtain histograms using Prism GraphPad.

### Immunofluorescence coupled with FISH (IF-FISH) for telomere dysfunction-induced foci (TIFs) analysis

1×10^5^ cells were seeded on coverslips and treated as indicated. Following treatment and recovery, cells were washed twice with PBS and fixed at room temperature with 4% formaldehyde (PFA) for 10 min. Fixed cells were rinsed with 1% BSA in PBS, and washed 3 times with PBS-Triton 0.2% before blocking with 10% normal goat serum, 1% BSA, and 0.1% Triton-x. Cells were incubated overnight at 4 °C with the indicated primary antibodies (anti-gammaH2AX (Ser139) mouse monoclonal, Santa Cruz #sc-517348, IF dilution 1:200; anti-53BP1 rabbit polyclonal, Novus #NB100-304, IF dilution 1:1000). The next day cells were washed with PBS-T 3 times before incubating with secondary antibodies and then washed again 3 times with PBS-T. Cells were then re-fixed with 4% formaldehyde and rinsed with 1% BSA in PBS, and then dehydrated with 70%, 90%, and 100% ethanol for 5 min. FISH was performed as described above, using the telomeric PNA probe TelC. Coverslips were mounted on slides with DAPI-containing mounting with ProLong™ Diamond Antifade Mountant (Thermo Fisher). Image acquisition was performed with a Nikon Ti inverted fluorescence microscope. Z stacks of 0.2 μm thickness (60x objective) were captured and images were deconvolved using NIS Elements Advance Research software algorithm. Telomere dysfunction-induced foci (TIFs) were defined as 53BP1 and γH2AX foci colocalizing with telomeres as described previously [4]. For micronuclei (MN) counting at least 300-400 nuclei were scored per experiment.

### Genomic DNA preparation for whole-genome sequencing

Genomic DNA was extracted using QIAGEN® Genomic DNA Preparation. Briefly, cells were collected by trypsinization 24h after the last exposure to telomeric 8-oxoG (N18), washed twice in cold PBS and resuspended in 0.5mL ice-cold PBS 1X. Cytoplasmic fraction was first extracted twice with buffer C1, and then the purified nuclear fraction was resuspended in buffer G2. 25 μL Proteinase K was added, and samples were incubated at 50°C for 1h. Both buffers contained the antioxidants butylated hydroxytoluene (BHT - Sigma) and deferoxamine mesylate salt (desferal - Sigma). DNA was then loaded on pre-equilibrated Genomic-tip 20/G with buffer QBT. The column was washed twice with buffer QC, and DNA was finally eluted using 2 mL of buffer QF. Genomic DNA was precipitated using 0.7 volumes isopropanol and resuspended in 200 μL 10mM Tris-HCl pH 8. Concentration and purity was measured by Nanodrop 3000 and 1 μg DNA per condition was provided for sequencing.

### DNA sequencing and analysis of telomere variants

Whole genome sequencing was obtained using paired-ended Illumina NovaSeq sequencing (Novogene). Telomere variant analysis was carried out using a previously described computational pipeline with minor modifications [22]. Briefly, sequencing reads enriched for telomeric content were identified using the GEAR software by selecting reads containing at least five consecutive telomeric repeats, including canonical (TTAGGG) and variant motifs, as well as their reverse complements. Filtered telomeric reads were aligned to the human reference genome (hg38) using minimap2 [23], and reads mapping to interstitial genomic regions with mapping quality >20 were excluded. For telomere variant analysis, telomeric reads were further mapped to a telomeric reference sequence, and the frequency of variant repeats was quantified using GEAR. Variant composition was determined based on deviations from the canonical TTAGGG repeat and reported as the relative abundance of specific telomeric sequence variants.

### Exo-FISH

Exo-FISH was performed as previously described [8,24,25]. Briefly, 1×10^6^ cells were seeded and then treated the next day. After recovery, cells were harvested and swollen in hypotonic solution (0.56% KCl), and fixed in methanol:acetic acid (3:1) prior to spreading onto glass slides. After rehydration, slides were treated with RNase A and incubated either in the absence or presence of exonuclease III (ExoIII, NEB). Slides were then dehydrated and hybridized with a Cy3-labeled telomeric (TTAGGG)₃ probe, followed by standard washes and mounting with DAPI-containing medium. Telomeric fluorescence intensity was quantified as a measure of exonuclease-accessible single-stranded DNA. For experiments involving chronic telomeric damage, cells were collected 3h after the final dye + light treatment (N18). Where indicated, slides were additionally incubated with formamidopyrimidine DNA glycosylase (FPG) prior to ExoIII digestion to convert 8-oxoG into SSBs. Specifically, following RNase A treatment, slides were washed and incubated with FPG in the appropriate buffer for 1h at 37 °C, washed again, and then processed for ExoIII digestion and telomeric probe hybridization as described above. For acquisition and analysis, images were taken throughout the slide avoiding aggregates to prevent signal intensity dependency on cell aggregation and/or uneven spread onto slides. Exposure time (ms) for both DAPI and Cy3 were kept consistent across experimental conditions in each separate experiment. Background signal was subtracted through blind iterative deconvolution (NIS Elements AR software). After conversion to Max-IPs, ROIs were defined through the DAPI channel, while relevant exo-FISH foci were selected setting a threshold for the Cy3 channel. Both lookup tables (LUTs) and the threshold were kept constant across experimental conditions throughout the analysis of an independent experiment. The automated measurement of the mean signal intensity for the thresholded channel (Cy3) was used on ROIs.

### S1-END-sequencing and END-sequencing

HeLa FAP-TRF1 WT and DKO cells were treated with dye and light as previously described, then returned to the incubator to recover for either 30 min or 24 h. Untreated cells were maintained in parallel as controls. S1-END-seq and END-seq were then performed as previously described [26–28]. Following treatment and recovery, cells were washed twice with PBS, then 10⁷ cells were embedded in 1% agarose plugs and incubated with proteinase K solution for 1 h at 50°C, followed by 7 h at 37°C. Plugs were then washed consecutively in wash buffer (10 mM Tris pH 8.0, 50 mM EDTA) and TE buffer (10 mM Tris pH 8.0, 1 mM EDTA), treated with RNase A (Puregene, QIAGEN), washed again in wash buffer, and stored at 4°C for up to two weeks. End blunting, A-tailing, and ligation to biotinylated hairpin adaptor 1 (ENDseq-adaptor-1, 5′-Phos-GATCGGAAGAGCGTCGTGTAGGGAAAGAGTGUU[Biotin-dT]U[Biotin-dT]UUACACTCTTTCCCTACACGACGCTCTTCCGATC∗T-3′;*phosphorothioate bond) were carried out in the plug to minimize shearing and in vitro damage. High molecular weight DNA was then isolated from the plugs and fragmented by sonication. Adaptor-ligated fragments were captured using streptavidin-coated beads, after which sonication-generated ends were repaired, A-tailed, and ligated to a distal adaptor (ENDseq-adaptor-2, 5′-Phos- GATCGGAAGAGCACACGTCUUUUUUUUAGACGTGTGCTCTTCCGATC∗T-3′;∗phosphorothioate bond). PCR amplification of the resulting material yielded sequencing-ready libraries in which the first sequenced base corresponds precisely to the first base of the blunted DSB on either side of the break. For S1-END-seq, proteinase K- and RNase A-treated plugs were incubated with 100 U of S1 nuclease (Thermo Fisher) at 37°C for 30 min. The reaction was quenched by addition of EDTA to a final concentration of 10 mM, and plugs were subsequently processed through the standard END-seq protocol described above. All libraries were sequenced on a NextSeq 2000 platform (Illumina) using 75-bp single-end read kits. For each treatment condition, control (untreated) and sample libraries were sequenced in parallel on the same flow cell. Telomeric sequences were identified and quantified from END-seq libraries using a custom Python script, getTelomereENDseq, run in a Linux command-line environment as described in [26], with a minimum of four consecutive CCCTAA and TTAGGG repeats required for C-rich and G-rich read classification, respectively.

### Statistics and reproducibility

The number of biological replicates is noted in all figure legends and methods. All statistical analyses were performed in GraphPad Prism 9 and 10. No statistical method was used to predetermine sample size. Investigators were not blinded to allocation during experiments and outcome assessments.

## RESULTS

### MUTYH deficiency increases chronic telomeric 8-oxoG induced telomere shortening and aberrations

To selectively generate 8-oxoG at telomeres, we used a previously described chemoptogenetic system in which a fluorogen-activating peptide (FAP) is fused to the telomeric protein TRF1. Upon binding of the photosensitizer di-iodinated malachite green (MG2I) dye and illumination with 660 nm light, the FAP-TRF1 complex produces singlet oxygen (^1^O₂). This dye + light (DL) treatment leads to localized telomeric 8-oxoG formation in HeLa cells [4,29,30]. To determine the role of MUTYH in the response to telomeric 8-oxoG damage, we used CRISPR/Cas9 and single cell cloning to generate knock out (KO) clones (Figure 1A). To confirm loss of MUTYH activity, we used the Fluorescence Multiplex based Host Cell Reactivation reporter (FM-HCR) assay, in which the 8-oxoG lesion is synthesized in the coding sequence of a reporter gene on a plasmid. Specifically, 8-oxoG paired with adenine in pMax_mPlum plasmid serves a substrate for MUTYH, while 8-oxoG paired with cytosine in pMax_mOrange plasmid is a substrate for OGG1 [19]. Unrepaired plasmids cause a transcriptional error that induces fluorescent reporter expression, thus, reporter expression is inversely proportional to repair capacity, whereas efficient repair results in low fluorescent reporter expression rate [19,21]. As expected, MUTYH KO cell lines failed to remove adenine from 8-oxoG:A, as indicated by significantly higher mPlum reporter expression compared to wild type (WT) and OGG1 KO cells [4,31], which are MUTYH-proficient (Figure 1B). Conversely, low mOrange reporter expression of the 8-oxoG:C lesion confirmed that MUTYH KO cells are proficient for repairing 8-oxoG:C lesions, in contrast to high expression observed in OGG1 KO cells (Figure 1C).

**Figure 1.**
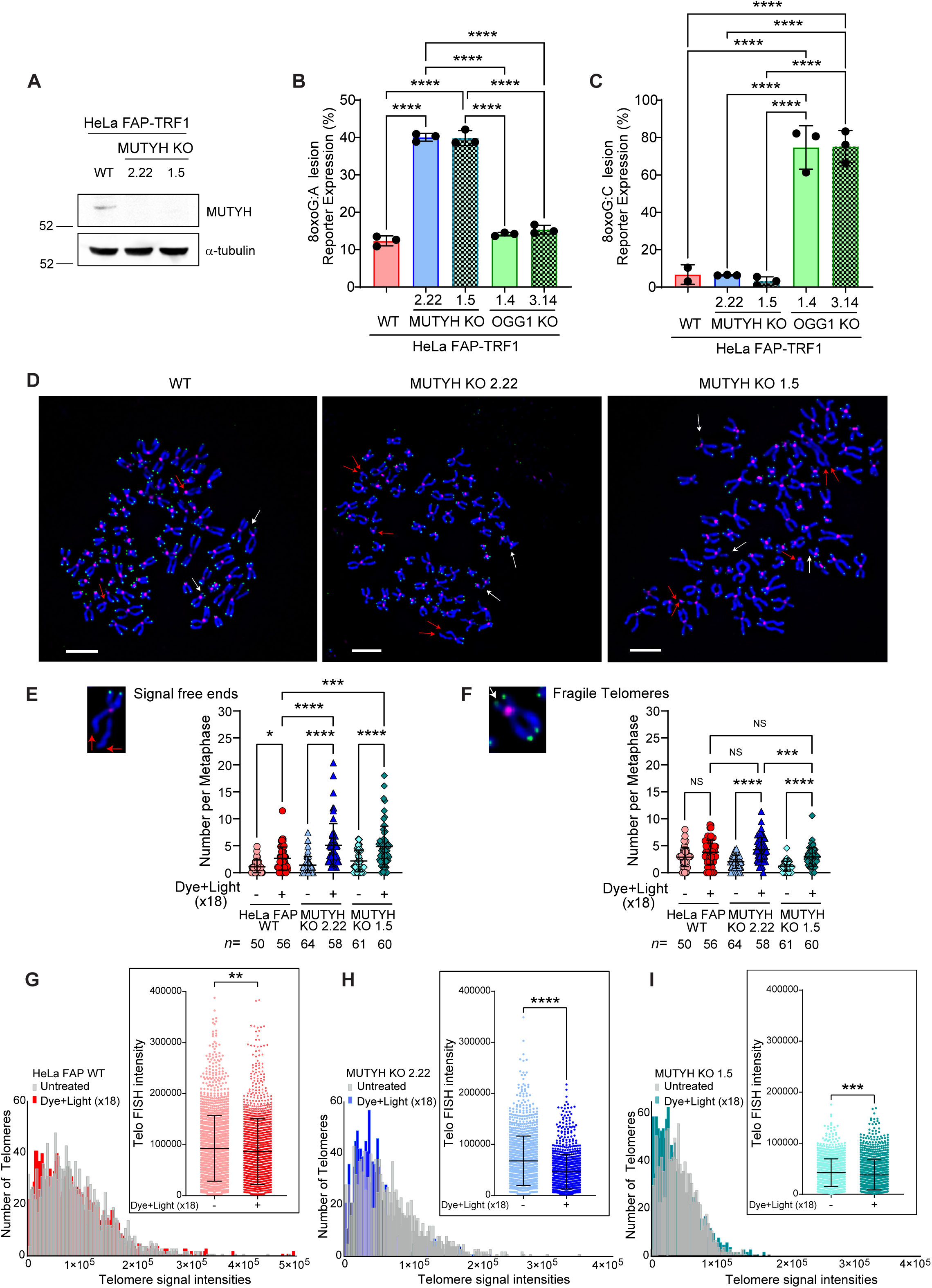
MUTYH deficiency increases chronic telomeric 8-oxoG induced telomere shortening and aberrations. **(A**) Immunoblot of MUTYH in parental HeLa FAP-TRF1 and two MUTYH KO individual clones. α-tubulin was used as a loading control. **(B-C)** FM-HCR was performed to assess DNA repair capacity in the indicated cell lines for BER reporters harboring a site specific (B) 8-oxoG:A or (C) 8-oxoG:C base lesion. Data represent the mean ± s.d. from three independent experiments. Statistical difference determined by one-way ANOVA with Tukey’s multiple comparisons; **** P<0.0001. Only comparisons yielding significant P-values are shown. **(D)** Telo-FISH of metaphase chromosomes 24 h after last dye + light (N18). Telomeric signal- free ends (red arrowheads) and fragile telomeres (white arrowheads) are indicated. Green = telomeres; pink = centromeres. Scale bars = 5 μm. **(E-F)** Number of telomeric signal-free chromatid ends (E) or fragile telomeres (F) per metaphase in the indicated cell lines. Data represents mean ± s.d. from the indicated n number of metaphases analyzed from three independent experiments, normalized to the chromosome number. Ordinary one-way ANOVA with Tukey’s multiple comparisons test; * P<0.05, ** P<0.01, *** P<0.001, **** P<0.0001. Insets show representative images of metaphase chromosomes scored as signal free ends or fragile by telo-FISH. **(G-I)** Quantification of telomeric signal intensities from telo-FISH of metaphase chromosomes (Figure 1F) from untreated and dye + light-treated HeLa FAP-TRF1 (G) WT, (H) MUTYH KO 2.22, and (I) MUTYH KO 1.5 cells. The x axis is shown as binning by 2,500 (a.u.). The inset shows the statistical unpaired t test on the single telomere FISH intensities; ** P<0.01, *** P<0.001, **** P<0.0001.

Having validated 8-oxoG:A repair deficiency in MUTYH KO cells, we next used our chemoptogenetic system to target 8-oxoG formation at telomeres. Exposing WT and MUTYH KO cells to 5 min DL to induce telomeric 8-oxoG formation did not increase telomeric aberrations (Supplementary Figure 1A-B) or telomere dysfunction-induced foci (TIFs) 24 h following damage (Supplementary Figure 1C-D). Similarly, we observed previously that OGG1 KO HeLa LT cells are also largely insensitive to acute telomeric 8-oxoG damage. However, chronic telomere damage induced more telomere aberrations and shortening in OGG1 KO cells compared to WT cells [4]. Therefore, to assess the importance of MUTYH activity in maintaining telomeres, we subjected WT and MUTYH KO HeLa FAP-TRF1 cells to repeated targeted 8-oxoG production, to mimic the conditions of chronic oxidative stress that cancer cells typically endure [4]. Cells were exposed to 5 min DL once per day over 24 days, except every 4^th^ day when cells were harvested for analysis, for a total of 18 treatments (Supplementary Figure 1E). While we did not observe a significant reduction in cell population doubling in response to damage in the MUTYH KO cells, unlike WT cells (Supplementary Figure 1F), analysis of metaphase chromosomes by telomere fluorescence in situ hybridization (telo-FISH) revealed that MUTYH deficiency caused a significantly higher number of telomere signal free ends after chronic telomeric 8-oxoG compared to WT cells (Figure 1D-E). In addition, chronic damage significantly increased fragile telomeres in MUTYH KO cell lines, but not in WT cells (Figure 1F). We then examined telomere length by quantitative telo-FISH and found that MUTYH KO cells showed a marked shift towards shortened telomeres after 18 exposures to DL compared to WT cells (Figure 1G-I). Taken together, these data show that MUTYH deficiency exacerbates telomere shortening and aberrations caused by chronic 8-oxoG damage, without significantly reducing cell growth.

### MUTYH loss increases genomic instability induced by chronic telomeric oxidative damage

Telomere losses can drive genomic instability by rendering chromosome ends susceptible to end-joining repair, generating dicentric chromosomes that mis-segregate during mitosis and ultimately produce micronuclei [32–34]. Because MUTYH KO cells exhibited increased signal-free ends after chronic telomeric 8-oxoG damage, we first asked whether apparent telomere losses corresponded to increased chromosomal end fusions. Following 18 DL treatments, MUTYH KO cells displayed a significantly higher percentage of metaphases containing one or more dicentric chromosomes compared with WT cells (Figure 2A-B). These aberrant chromosomes often arise when dysfunctional telomeres from different chromosomes are recognized as DSBs and repaired through aberrant end-joining [35]. To determine whether persistent telomeric DNA damage signaling accompanied chronic 8-oxoG formation, we quantified TIFs. However, we observed no significant increase in colocalization of telomeres with the DNA damage response (DDR) markers γH2AX and 53BP1 in any of the cell lines after chronic DL treatments (Figure 2C-D), indicating that the genomic instability in MUTYH KO cells does not stem from persistent telomeric DDR activation. Since dicentric chromosomes often break during anaphase and produce chromosome fragments that can be encapsulated into micronuclei [36], we next assessed micronuclei formation over the course of chronic 8-oxoG exposure. MUTYH KO cells exhibited elevated micronuclei after as early as 9 treatments, and this difference became more pronounced after 18 treatments, with both MUTYH-deficient clones displaying significantly more micronuclei than WT (Figure 2E-G). Together, these findings demonstrate that chronic telomeric 8-oxoG damage promotes greater genomic instability in MUTYH-deficient cancer cells, compared to WT, manifested by increased dicentric chromosome formation and micronuclei.

**Figure 2.**
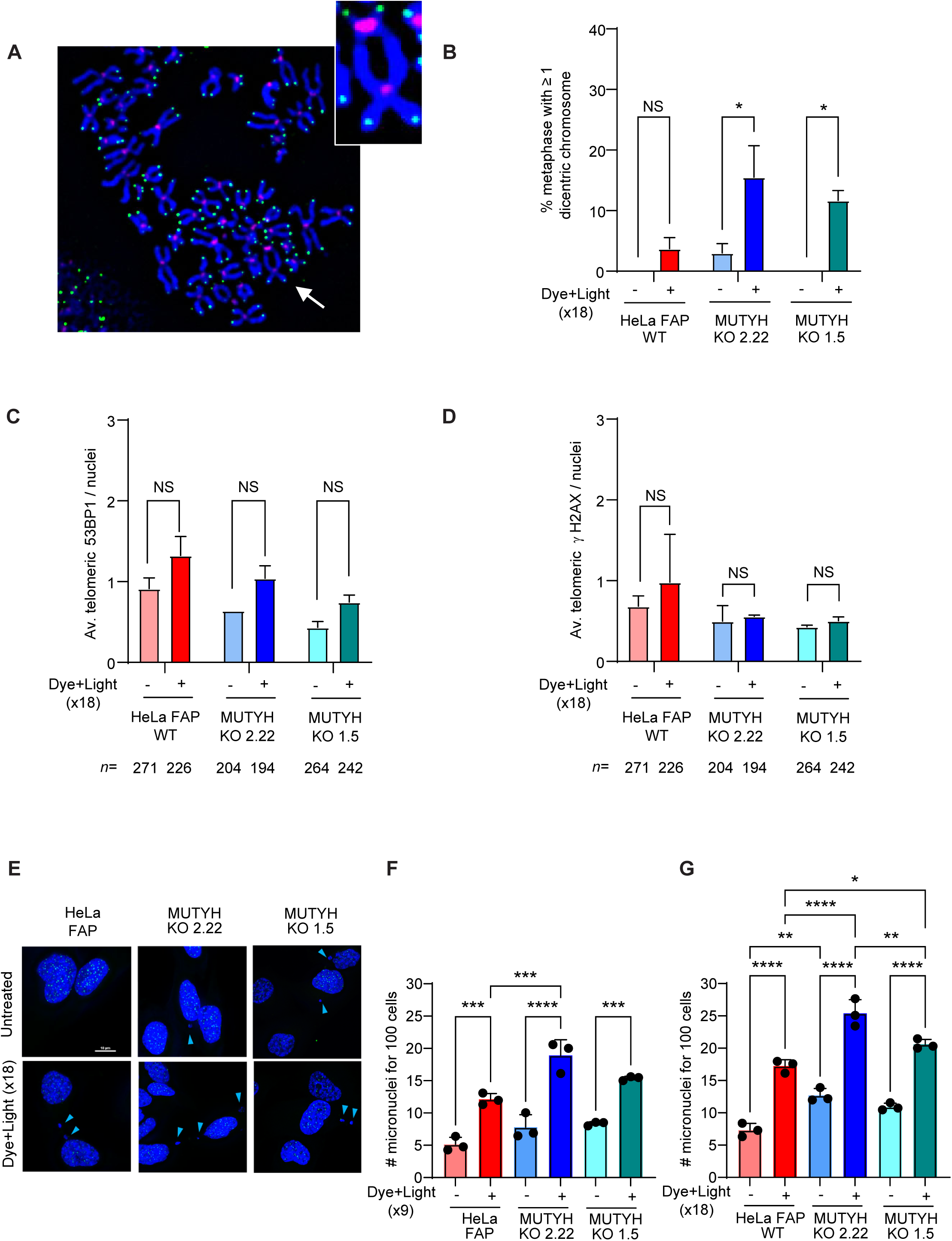
MUTYH loss increases genomic instability induced by chronic telomeric oxidative damage. **(A)** Telomeric (green) and centromeric (pink) staining by FISH on metaphase chromosomes 24 h after treatment 18. A dicentric chromosome is indicated by a white arrow. **(B)** Percentage of metaphases with at least one dicentric chromosome. Means ± SEM from 3 independent experiments (same number of metaphases analyzed as in Figure 1E-F); * P<0.05, two-way ANOVA. **(C-D)** Quantification of average (C) telomeric 53BP1 and (D) telomeric γH2AX per nucleus after 18 treatments. Means ± SEM from the indicated *n* number of nuclei analyzed from two independent experiments; one-way ANOVA, ns = non significant. **(E)** Micronuclei in HeLa FAP-TRF1 cells after 18 treatments. DNA is stained by DAPI and telomeres are stained with green fluorescent PNA probes. Blue arrows point to micronuclei. **(F-G)** Quantification of micronuclei per 100 cells after treatments 9 (F) and 18 (G). Means ± s.d. from 3 independent experiments (at least 300 nuclei per condition for each experiment). * P<0.05, ** P<0.01, *** P<0.001, **** P<0.0001, one-way ANOVA.

### Chronic telomeric 8-oxoG damage in MUTYH KO cells increases G to T mutations

Given MUTYH’s role in suppressing mutations and our result that MUTYH loss increases telomere instability from chronic telomeric 8-oxoG damage, we next asked whether chronic lesions also leave a detectable mutational footprint within the telomeric sequence. Unrepaired 8- oxoG lesions can mispair with adenine and generate G:C→T:A transversion mutations [37]. To determine whether chronic telomeric 8-oxoG leads to the accumulation of telomere sequence variants, we performed whole-genome sequencing (WGS) on DNA isolated from WT and MUTYH KO cells subjected to 18 rounds of DL treatment. Telomere-enriched reads (≥6 consecutive TTAGGG) were extracted and enumerated to detect variant telomeric repeats, which were normalized to total TTAGGG repeats. Consistent with the expected unrepaired A:8-oxoG mispairs, DL-treated MUTYH KO cells exhibited a detectable increase in variant telomere sequences compared with both untreated MUTYH KO cells and DL-treated WT controls. In particular, while WT cells displayed only minimal changes in variant composition, variants consistent with G→T transversions, including TTATGT, TTATTG, and TTATTT, were enriched following chronic 8-oxoG exposure in the MUTYH-deficient background (Figure 3A-B). These findings indicate that MUTYH activity is required to prevent 8-oxoG-mediated telomere mutagenesis.

**Figure 3.**
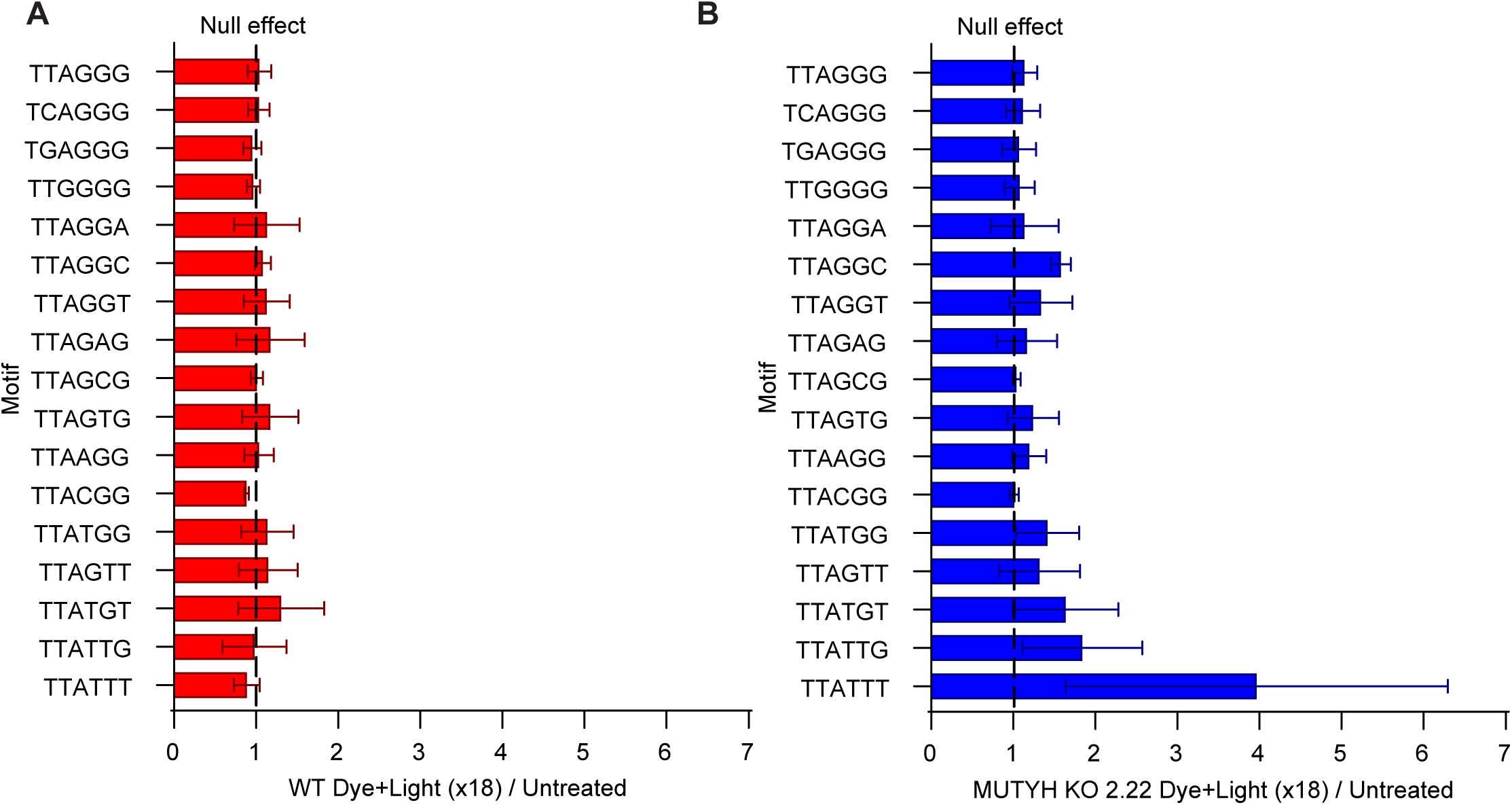
Chronic telomeric 8-oxoG damage in MUTYH KO cells increases G to T mutations. **(A-B)** Normalized and proportional count of telomere variant repeats relative to TTAGGG repeats. The ratio of variant telomere repeats to TTAGGG repeats is shown. Telomere variant repeats whose increase in MUTYH KO cells is proportionally greater than that of the TTAGGG repeat exceed the dotted line. Error bars refer to the range of values from three independent experiments.

### Loss of both MUTYH and OGG1 glycosylases rescues chronic damage-induced telomere aberrations and genomic instability

Our data show that MUTYH activity is crucial for maintaining telomere and genomic stability in response to persistent telomeric 8-oxoG damage, similar to our prior results with OGG1 [4]. Since both glycosylases work to prevent the detrimental effects of oxidative damage at telomeres, we next examined the consequences of telomeric 8-oxoG in cells doubly deficient for MUTYH and OGG1. For this, we knocked out both glycosylases in the HeLa FAP-TRF1 cells and obtained individual double knock out (DKO) clones (Figure 4A), which we confirmed were deficient for both removal of A opposite 8-oxoG and 8-oxoG opposite C repair (Figure 4B-C). Chronic telomeric 8-oxoG damage did not significantly reduce the population doubling of DKO cells (Supplementary Figure 2A). Surprisingly, analysis of telomere aberrations on metaphase chromosomes revealed that the increased damage-induced telomere losses and fragility caused by MUTYH KO (Figure 1E-F) and OGG1 KO [4], compared to WT cells, were rescued in the DKO cells (Figure 4D-E). Telomere length analysis by quantitative telo-FISH revealed that DKO cells had significantly longer telomeres after 18 exposures to DL (Figure 4G-H), in contrast to WT and MUTYH KO cells, which experienced telomere shortening (Figure 4F and 1G-I), and to OGG1 KO cells, as our previous data had shown [4]. Chronic telomere damage did not significantly increase the number of dicentric chromosomes in DKO cells (no dicentrics observed, data not shown), and the percentage of micronuclei produced after chronic oxidative damage in DKO cells was similar to WT cells (Figure 4I). Taken together, these results suggest that the activity of MUTYH and OGG1 glycosylases in the context of dysregulated BER, when one glycosylase is deficient, is more detrimental than leaving 8-oxoG unrepaired.

**Figure 4.**
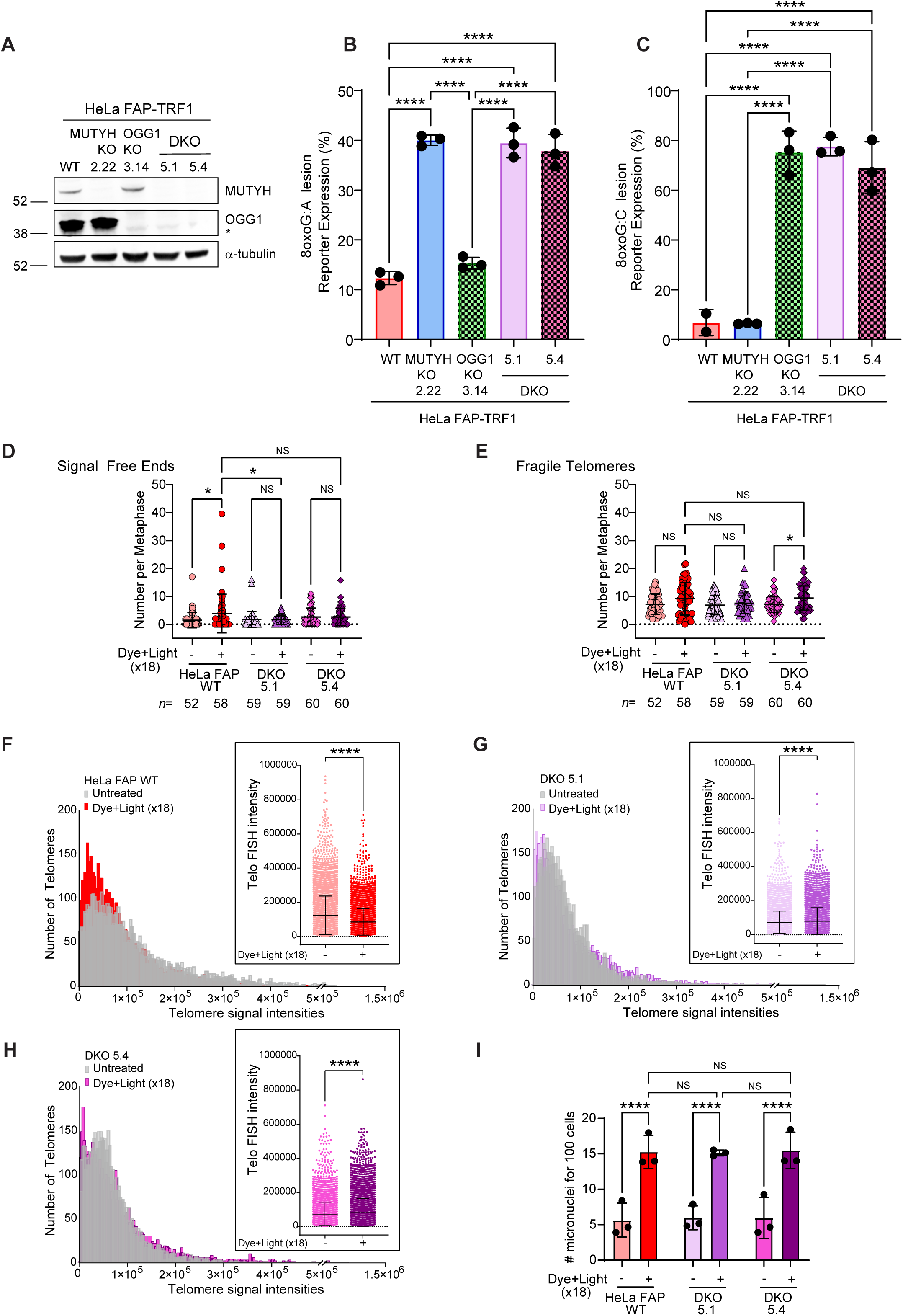
Loss of both MUTYH and OGG1 glycosylases rescues chronic damage-induced telomere aberrations and genomic instability. **(A**) Immunoblot of MUTYH and OGG1 in parental HeLa FAP-TRF1 and MUTYH KO, OGG1 KO or DKO individual clones. α-tubulin was used as a loading control. * indicates nonspecific band below OGG1. **(B-C)** FM-HCR assay was performed to assess DNA repair capacity in the indicated cell lines for BER reporters harboring a site specific (B) 8-oxoG:A or (C) 8-oxoG:C base lesion. Data are normalized to WT cells and represent the mean ± s.d. from three independent experiments. Statistical difference determined by one-way ANOVA with Tukey’s multiple comparisons; **** P<0.0001. **(D-E)** Number of telomeric signal-free chromatid ends (D) or fragile telomeres (E) per metaphase in the indicated cell lines. Data represents mean ± s.d. from the indicated *n* number of metaphases analyzed from three independent experiments, normalized to the chromosome number. Ordinary one-way ANOVA with Tukey’s multiple comparisons test; ns=non significant, * P<0.05. **(F-H)** Quantification of telomeric signal intensities from telo-FISH of metaphase chromosomes from untreated and dye + light-treated HeLa FAP-TRF1 (F) WT, (G) DKO 5.1, and (H) DKO 5.4 cells. The x axis is shown as binning by 2,500 (a.u.). The inset shows the statistical unpaired t test on the single telomere FISH intensities; **** P<0.0001. **(I)** Quantification of micronuclei per 100 cells after 18 treatments. Means ± s.d. from three independent experiments (at least 300-400 nuclei per condition for each experiment); ns = non significant, *** P<0.001, **** P<0.0001, 1-way ANOVA.

### Glycosylase deficiency suppresses BER-induced SSBs after chronic damage

Both MUTYH and OGG1 produce SSB repair intermediates during BER of misincorporated adenines or 8-oxoG, respectively, and we previously showed that SSBs are rapidly detected at telomeres 30 min after targeted 8-oxoG induction in non-diseased cells [8,38]. The accumulation of SSB intermediates suggests that BER completion at telomeres may be less efficient in these cells. Therefore, we asked if 8-oxoG BER produces SSB intermediates in HeLa LT cancer cells, since this could explain why the loss of both glycosylases suppresses telomere instability after damage. For this, we used the exo-FISH method. In this assay, exonuclease III activity at SSBs produces single-stranded telomeric DNA to which a fluorescent (TTAGGG)_3_ telomeric probe is annealed and quantified. Fluorescence signal intensity is directly proportional to the amount of SSBs present at telomeres [8,24,25]. Furthermore, to determine if unrepaired 8-oxoG lesions accumulate in DKO cells, we also pretreated slides with the bifunctional glycosylase formamidopyrimidine DNA glycosylase (FPG), which removes unrepaired 8-oxoG lesions and cleaves the DNA backbone [8]. The difference in signal between +EXO alone and +FPG +EXO together indicates 8-oxoG lesions.

We repeatedly treated WT, MUTYH KO and DKO cells with DL for chronic telomeric 8-oxoG induction, and measured SSBs and 8-oxoG lesions 3 h after the last treatment (N18). We reasoned that by 3 h, most of the 8-oxoG lesions should be repaired in WT cells based on OGG1 kinetics and departure from telomeres at this recovery time point [4,31]. Surprisingly, WT and MUTYH KO cells showed a significant increase in SSB intermediates even after 3 h recovery (Figure 5A- B, + EXO). No further increase upon FPG addition indicated a lack of persistent 8-oxoGs in WT cells. However, the + FPG +EXO signal was significantly higher in MUTYH KO compared to WT, suggesting delayed 8-oxoG repair. In contrast, DKO cells showed a significant increase in +FPG +EXO signal, compared to +EXO alone, prior to damage, which increased further after DL treatment. DL also increased the EXO signal slightly in DKO cells, indicative of SSBs, although to a lesser extent than in WT and MUTYH KO cells. Taken together, increased SSBs in WT and MUTYH KO cells are consistent with incomplete 8-oxoG BER at telomeres, and the accumulation of 8-oxoGs in glycosylase deficient cells confirms a lack of repair.

**Figure 5.**
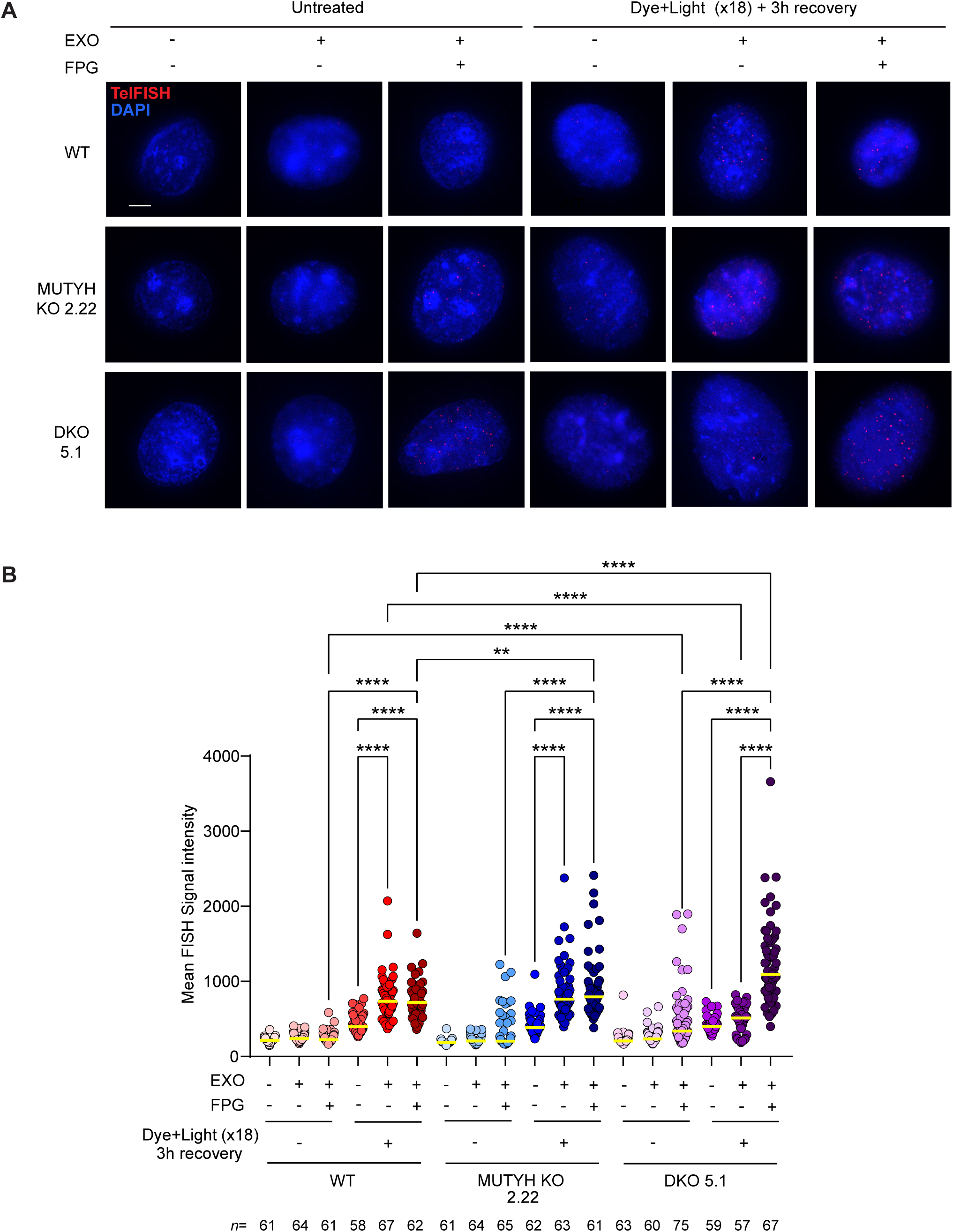
Glycosylase deficiency suppresses BER-induced single strand breaks after chronic damage. **(A)** Representative images of exo-FISH performed in cells untreated or treated 18 times with 5 min dye + light and harvested 3 h after the last treatment, showing telomeres harboring SSBs on the G-rich strand (red foci). Cells were pre-treated with or without FPG and exonuclease III as indicated, before hybridization with TelG FISH probe. Scale bar = 5 μm. **(B)** Quantification of exo-FISH signal intensity in cells untreated or treated with dye + light for 18 times and recovered 3 h before being fixed. Cells were treated with FPG or exonuclease III, or both as indicated. Each data point represents the mean fluorescence intensity for each cell. Data represent the median of the indicated *n* number of nuclei analyzed per each condition from two independent experiments; ns = non significant, ** P<0.01, *** P<0.001, **** P<0.0001, two-way ANOVA.

### Doubly glycosylase deficient cells exhibit delayed SSB formation after telomere damage

We were surprised that the doubly glycosylase deficient cells showed a slight, but significant, increase in SSBs at 3 h recovery from the last treatment following chronic damage, since these cells should not initiate BER at 8-oxoG lesions. Therefore, we examined SSBs in WT and DKO cells shortly after damage (30 min) to allow time for BER initiation, and after 24 h to allow ample time for BER completion, following acute telomere damage. 20 min DL followed by 30 min recovery significantly increased the exo-FISH signal intensity in WT HeLa cells but not in DKO, as expected (Figures 6A-B), and we confirmed this increase was specific with controls lacking exonuclease treatment (Supplementary Figure 2B). Surprisingly, we detected a significant increase in exo-FISH signal intensity in the DKO cells 24 h after the damage, whereas WT showed few, if any, detectable SSBs at that time point, consistent with BER completion after such long recovery (Figure 6B).

**Figure 6.**
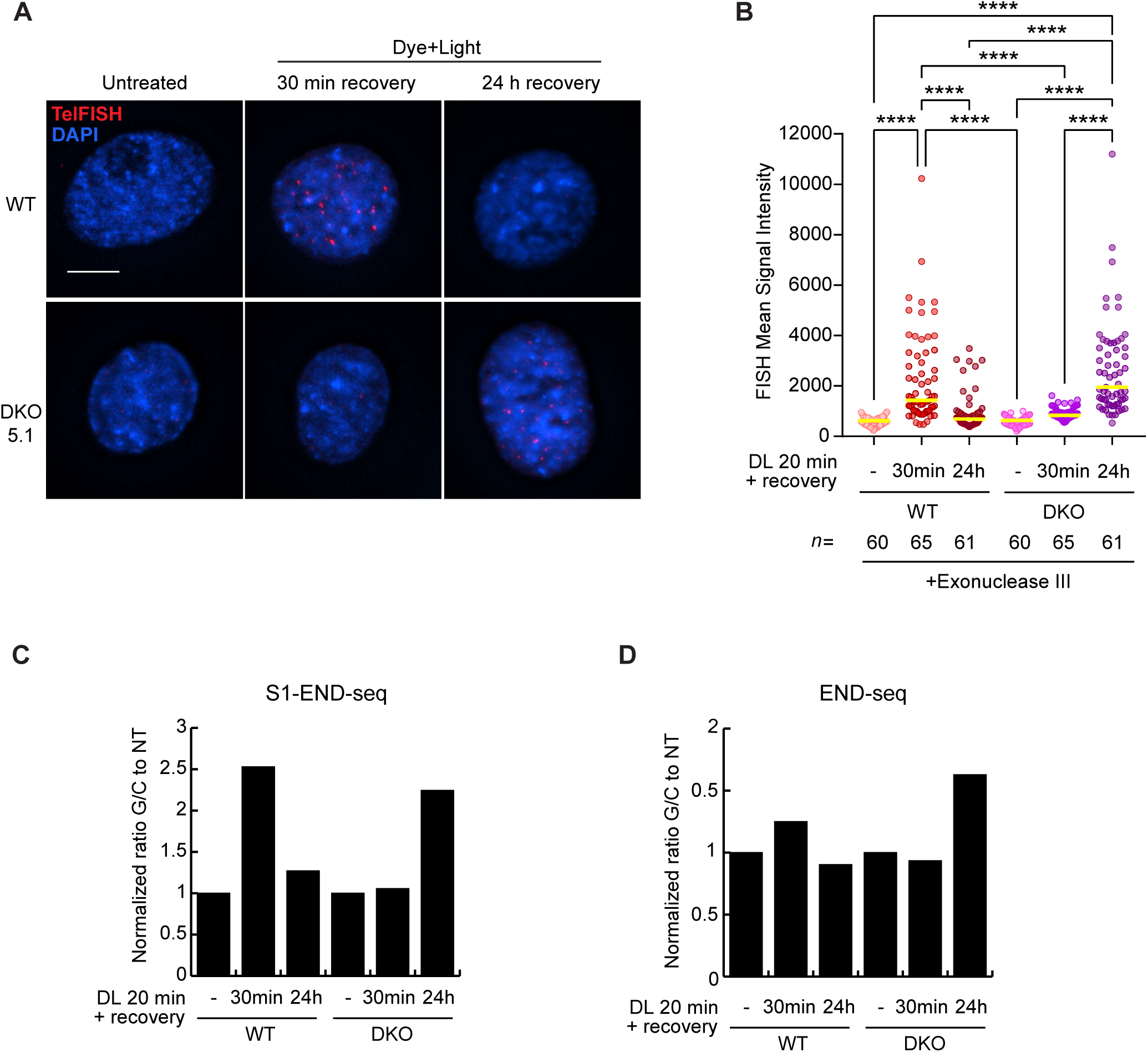
Targeted telomeric 8-oxoG induces rapid formation of SSBs intermediates in WT cells and DSBs only in repair deficient cells. **(A)** Representative images of exo-FISH performed in cells untreated, treated with 20 min dye + light and recovered for the 30 min or 24 h, showing telomeres harboring SSBs on the G-rich strand (red foci). Cells were treated with exonuclease III before hybridization with TelG FISH probe. Scale bar = 10 μm. **(B)** Quantification of exo-FISH signal intensity from Panel A. Each data point represents the mean fluorescence intensity value for each nucleus. Data represents the median of the indicated *n* number of nuclei analyzed from three independent experiments; **** P<0.0001, ordinary one-way ANOVA. **(C-D)** Normalized ratio of G-rich to C-rich telomeric reads (G/C) obtained by S1-END-seq (C) and END-seq (D) in untreated and dye + light treated cells recovered for the indicated times. Values are expressed relative to untreated controls. An increase in the G/C ratio reflects enrichment of S1-sensitive single-stranded regions within telomeres, indicative of accumulated repair intermediates (C), whereas in END-seq it reflects enrichment of double-strand breaks (D).

Since exo-FISH can also detect DSBs, we could not exclude that the significant increase in exo- FISH signal 24 h after damage in the DKO cell was also due to DSBs. While we previously demonstrated that the repair of telomeric 8-oxoG drives the production of SSB intermediates, and not DSBs, in non-diseased cells, this analysis was done only 30 min after damage [8]. To further distinguish between SSB and DSB formation, and to confirm our exo-FISH data, we carried out S1-END-seq, a modification of the high-resolution and unbiased sequencing method END-seq in which S1 nuclease treatment precedes the standard END-seq protocol [26,28]. In this assay, S1 nuclease cleaves single-stranded DNA, converting SSBs into two-ended DSBs that are subsequently captured and sequenced. In the absence of ssDNA, standard END-seq detects telomeres as one-ended DSBs. In contrast, S1-mediated cleavage of ssDNA within the internal telomere tract generates two-ended DSBs. [26–28]. Consistent with our exo-FISH results, S1-END-seq revealed 2.5 times more S1-dependent telomeric cleavage events in WT cells and no detectable signal in the DKO cells 30 min after 8-oxoG induction, compared to non-treated controls (Figure 6C, and Supplementary Figure 2C). However, 24 h post-damage, only the DKO cells showed about a 2-fold increase in S1-dependent cleavage, and slightly more DSBs as shown by standard END-seq (Figure 6D). When compared to the cancer cell line U2OS, which uses the alternative lengthening of telomeres (ALT) pathway and naturally displays elevated single-stranded and double-stranded damage at the telomeres [26], the amount of both DSBs and SSBs detected in HeLa LT cells appears generally smaller (Supplementary Figure 2D-E). This is consistent with the expected difference in the amount of telomeric breaks between telomerase-positive and ALT-positive cells [39]. In summary, these results indicate that BER-generated SSB intermediates in WT cells are eventually resolved, while glycosylase deficient cells exhibit a delayed appearance of telomeric breaks, 24 h post damage induction.

## DISCUSSION

Chronic oxidative stress is a pervasive feature of cancer cells, due to the elevated ROS produced by increased metabolism, hypoxia and exposure to chemotherapeutic agents [40–42], and is a major source of oxidative DNA damage and telomere dysfunction [43]. Among the repair proteins involved in 8-oxoG processing, OGG1 has an established role in reducing telomere attrition and crisis in cancer cells caused by oxidative damage [4,7,44]. Conversely, how MUTYH contributes to telomere protection in cancer cells under conditions of persistent oxidative damage has remained unclear. Here, by repeatedly inducing telomere-specific 8-oxoG lesions with our chemoptogenetic tool to mimic chronic oxidative stress, we uncovered a previously unrecognized requirement for MUTYH in maintaining telomere stability, preventing genome instability, and mitigating the burden of telomeric 8-oxoG damage in cancer cells.

Our findings on telomere attrition in MUTYH deficient cancer cells in response to chronic oxidative damage are consistent with a model in which the accumulation of unrepaired adenines misincorporated opposite 8-oxoG causes lesion persistence and increased mutagenesis. OGG1 cannot remove 8-oxoG when paired with adenine [45–47], and this might explain why Mutyh -/- mice have higher levels of genomic 8-oxoG following treatment with oxidants despite being proficient for OGG1 repair activity [48]. Consistent with this, MUTYH KO cells displayed increased unrepaired 8-oxoG lesions relative to WT 3 h after the last chronic exposure, suggesting delayed or incomplete repair of telomeric 8-oxoG in the absence of MUTYH. In the absence of a functional MUTYH, the persisting 8-oxoG at telomeres might trigger replication stress and promote telomere shortening and loss, similarly to our previous evidence in OGG1 deficient cells [4]. Despite experiencing increased telomere instability, MUTYH KO cells did not show significantly decreased proliferation following chronic 8-oxoG damage. This is consistent with data from Mutyh -/- mice and MAP patients, which have an elevated risk of spontaneous tumorigenesis, specifically colorectal cancer, and thus uncontrolled cancer cell proliferation [37,49].

Another mechanism by which telomere shortening and loss might arise following oxidative damage in MUTYH-deficient cancer cells is consequent telomere mutations impairing shelterin proteins, which bind TTAGGG repeat sequences to protect the telomeres [33]. Our sequencing analysis revealed that chronic telomeric 8-oxoG damage leads to a greater accumulation of telomere variants in MUTYH-deficient cells, including several variants consistent with G→T transversions. These findings suggest that the inability to excise adenines misincorporated opposite 8-oxoG allows them to become fixed as heritable alterations within telomeric DNA. Telomere variant repeats have been implicated in altered telomere-protein interactions, changes in telomere architecture, and increased recombination potential [50]. Notably, we did not detect sustained telomeric DDR signaling during chronic 8-oxoG exposure, indicating that the observed genomic instability likely arises from failure in repairing 8-oxoG rather than persistent damage signaling. To our knowledge, no clinical studies have examined telomere sequence composition in MAP patients or cancers with sporadic MUTYH mutations or decreased expression. Large-scale datasets such as COSMIC provide a comprehensive catalog of somatic mutations across many tumor types, including signatures specific to defective BER due to MUTYH mutations, such as SBS18 and SBS36 [51–53]. However, since telomeres are particularly difficult to sequence due to their highly repetitive nature, these datasets have not been able to capture telomeric repeat variants so far. Future work will be needed to determine the extent of telomere sequence mutagenesis in MUTYH-deficient human tumors. Nevertheless, our results provide the first experimental evidence that chronic oxidative damage can alter the telomeric DNA composition when BER of 8-oxoG is defective.

Our work has revealed that the detrimental consequences of chronic telomeric oxidative damage in cancer cells cannot be solely attributed to the presence of 8-oxoG itself, but also arise from dysregulated repair. The DKO cells lacking both OGG1 and MUTYH displayed a significant rescue of damage-induced telomere instability phenotypes, including losses and fragility, and micronuclei formation. These results point to BER intermediates, rather than 8-oxoG lesions, as key drivers of telomere dysfunction. The observation that DKO cells maintain telomere integrity despite accumulating unrepaired 8-oxoG underscores the vulnerability of telomeres to strand- incision steps during BER, consistent with our prior studies showing that SSBs and nicks pose challenges to telomeric replication [3,8]. Our exo-FISH and S1-END-seq analyses further support this model. WT cells displayed SSB intermediates on the G-rich strand shortly after 8-oxoG induction, consistent with rapid OGG1-mediated BER initiation, which persisted after 3 h recovery, but were resolved within 24 h. In contrast, DKO cells showed little evidence of SSB formation shortly after damage, but accumulated SSBs 24 h later. We speculate SSBs in DKO cells may have arisen from the excision of further 8-oxoG oxidation products, such as hydantoin lesions. These lesions are recognized and repaired by the NEIL1 and NEIL3 glycosylases, which can excise them even when embedded into telomeric G-quadruplex structures [54,55]. Although we did not find evidence for NEIL1 recruitment in WT FAP-TRF1 HeLa LT cells previously [4], we cannot exclude the possibility that NEIL glycosylases may be recruited when OGG1 and MUTYH are absent. Evidence from studies in MEFs suggests that another potential mechanism for the delayed appearance of SSBs in DKO cells may involve the removal of 8-oxoG opposite A by the mismatch repair (MMR) protein Msh2 in the newly synthesized DNA [56–58]. If MSH2 recognizes the 8- oxoG:A or 8-oxoG:C, and excises 8-oxoG, this would generate a single stranded DNA in the G-rich strand. However, since biochemical studies show 8-oxoG:C and 8-oxoG:A are poor substrates for MMR [59], this may explain the delayed formation of SSBs in DKO cells. The slight increase in DSBs observed in DKO cells 24 h after damage may have arisen from replication fork collapse [60], but they were clearly rare events.

Our result that OGG1 and MUTYH doubly deficient HeLa LT cells exhibit telomere elongation after chronic telomere damage was surprising, given the result that these cells accumulate telomeric 8-oxoG lesions. Our prior study revealed that OGG1 loss in HeLa LT cells caused persistent telomeric 8-oxoG lesions and exacerbated telomere shortening after chronic telomere damage [4], and the current study revealed a similar result of shortened telomeres in MUTYH KO cells after chronic damage. We also observed telomere elongation in non-diseased BJ hTERT fibroblasts deficient for both MUTYH and OGG1 after chronic telomere damage (unpublished data), suggesting that this effect is not restricted to cancer cells. Although the molecular basis of this telomere elongation remains to be determined, several mechanisms may contribute. Persistent 8-oxoG lesions can induce replication stress at telomeres, leading to fork stalling and collapse, which in the absence of both MUTYH and OGG1 may trigger recombination-based telomere maintenance pathways such as break-induced replication (BIR) [60,61]. In glycosylase deficient cells, unrepaired oxidative lesions may therefore promote engagement of ALT-like mechanisms, resulting in telomere extension. We observed previously that telomeric 8-oxoG can stimulate homologous recombination (HR) and ALT activity in ALT cancer cell lines, but not WT HeLa LT cells [62]. However, the appearance of SSBs and DSBs in HeLa LT DKO cells, but not WT, following 24 h recovery from damage and time for DNA replication, raises the probability for engagement of HR mechanisms in DKO cells to resolve the breaks. Together, these possibilities suggest that, in the absence of BER, chronic oxidative damage may shift the balance from telomere shortening toward aberrant elongation through alternative DNA repair-mediated mechanisms.

## CONCLUSION

Collectively, this study establishes MUTYH as a critical guardian of telomere integrity under chronic oxidative stress in cancer cells. MUTYH germline mutations in MAP-related cancers, and somatic mutations or deficiencies in sporadic cancers, contribute to tumorigenesis partly by increasing oxidative damage-induced mutagenesis in proto-oncogenes and tumor suppressor genes [8]. Beyond this established role, our results suggest MUTYH defects may also contribute to tumorigenesis by promoting senescence evasion [8] and genomic instability in response to oxidative telomeric damage. Our study further highlights the susceptibility of telomeres to repair intermediates and how incomplete or dysregulated base excision repair is a major source of telomere instability in cancer cells.

## AUTHOR CONTRIBUTIONS

M.D.R. and P.L.O. conceived the study and designed the experiments. M.D.R. performed most of the experiments. T.H. and L.C. assisted with telo-FISH experiments. M.D.R. and P.L.O. wrote the manuscript with assistance from the other authors. SMT: coordinated, designed and conducted the FM-HCR experimental work, data analysis & interpretation. ZDN: supervision & FM-HCR data interpretation. P.G., N.A. and H.A.P. analyzed the WGS data for the telomere mutagenesis analysis. B.A. and E.L.D. conducted END-seq and S1-END-seq and analysis.

## DATA AVAILABILITY STATEMENT

All data generated or analyzed during this study are included in this published article and its supplementary information files. The datasets used during the current study are available from the corresponding author on request. Further information and requests for reagents should be directed to and will be fulfilled by the corresponding author.

## ACKNOWLEDGEMENTS

We are grateful to Brigitte Schmidt and Bruce Armitage (Carnegie Mellon University) for providing MG2I dye.

## FIGURE LEGENDS

**Supplementary Figure 1.**
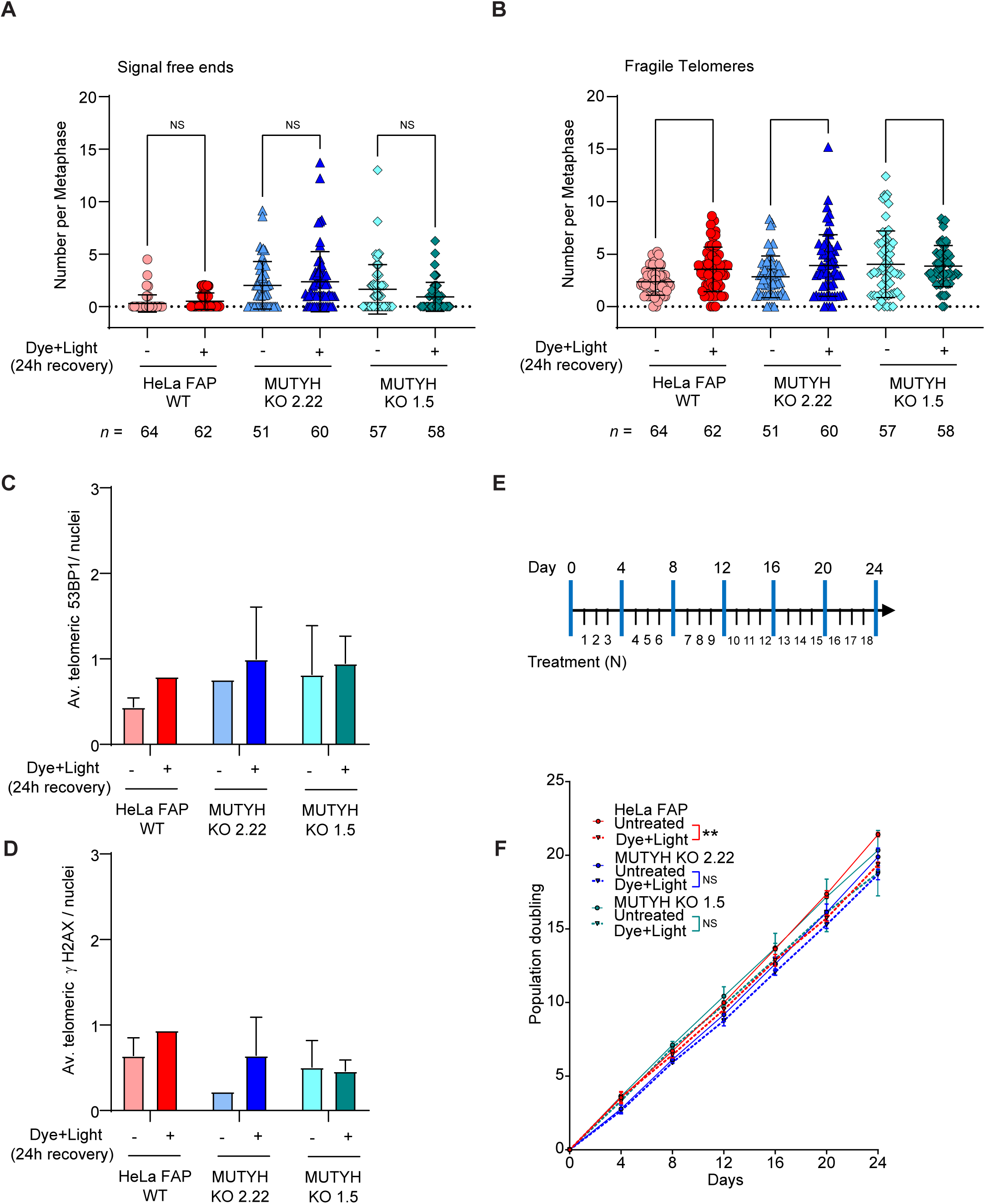
**(A-B)** Number of telomeric signal-free chromatid ends (A) or fragile telomeres (B) per metaphase in the indicated cell lines, untreated or treated with dye + light and recovered 24 h (acute exposure to telomeric 8-oxoG). Data represents mean ± s.d. from the indicated n number of metaphases analyzed from three independent experiments, normalized to the chromosome number. Ordinary one-way ANOVA; ns=non significant. **(C-D)** Quantification of average telomeric 53BP1(C) and telomeric γH2AX (D) per nucleus 24 h after treatment with dye + light. Means ± SEM from two independent experiments; one-way ANOVA, ns = non significant. **(E)** Schematic of the repeated telomeric 8-oxoG inductions. Blue bars indicate days when cells were harvested and not exposed. **(F)** Population doubling (PD) over 24 days of untreated cells (solid line) and cells treated with dye + light each day except every 4th day of harvest (dotted line). Data represents mean ± SEM from three independent experiments. Welch’s two-sample t test comparing untreated and Dye+Light at N18 per each cell line; ** P<0.005.

**Supplementary Figure 2.**
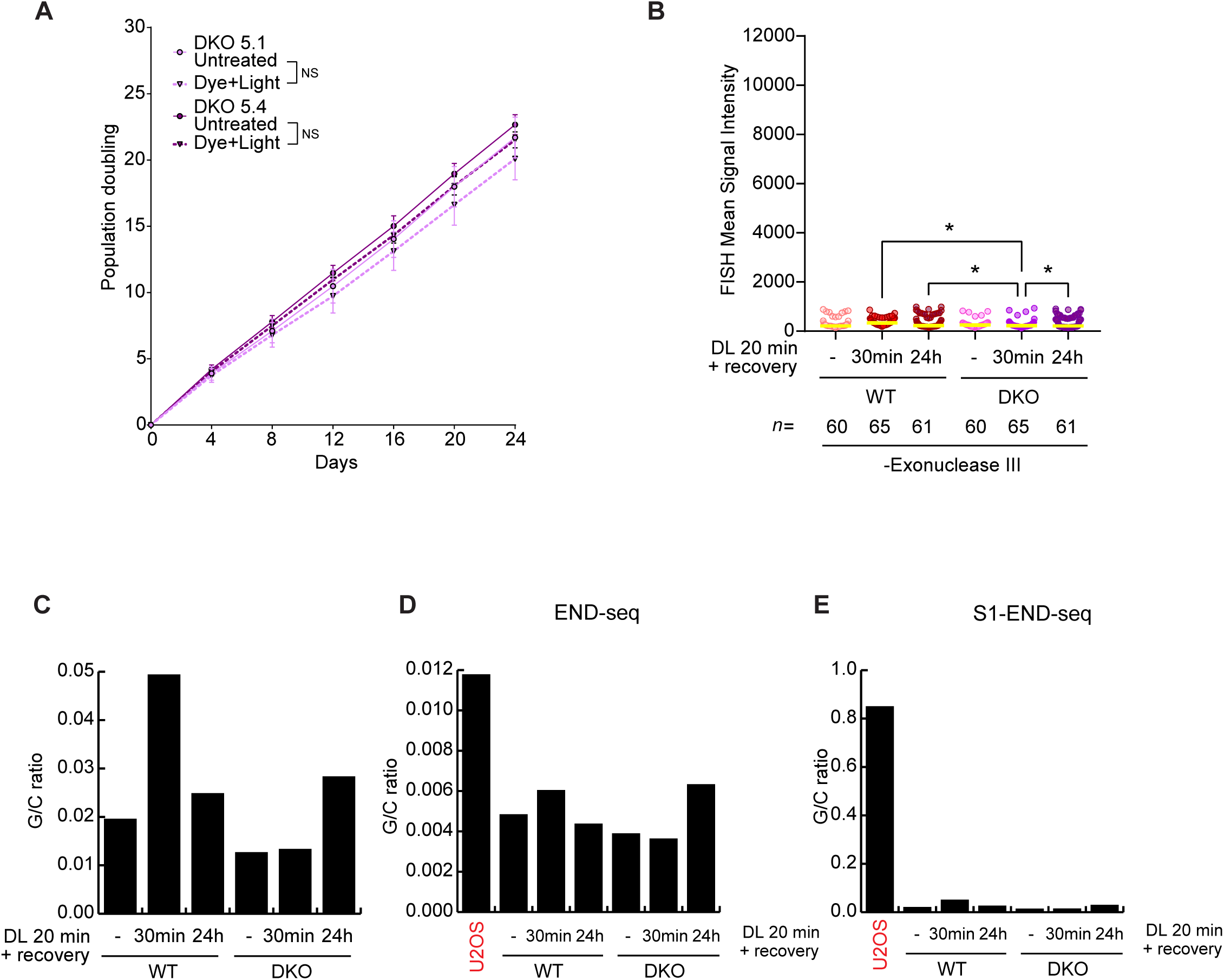
**(A** PD over 24 days of untreated cells (solid line) and cells treated with dye + light each day except every 4th day of harvest (dotted line). Data represents mean ± SEM from three independent experiments. **(B)** Quantification of exo-FISH signal intensity from the controls of panels in Figure 5A-B, without exonuclease III treatement. Each data point represents the mean fluorescence intensity value for each nucleus. Data represents the median of the indicated *n* number of nuclei analyzed from three independent experiments; * P<0.05, ordinary one-way ANOVA. **(C)** Ratio of G-rich to C-rich telomeric reads (G/C), not normalized on the untreated samples, obtained by S1-END-seq in untreated and dye + light treated cells recovered for the indicated times. An increase in the G/C ratio reflects enrichment of S1-sensitive single-stranded regions within telomeres, indicative of accumulated repair intermediates. **(D-E)** Ratio of G-rich to C-rich telomeric reads (G/C), not normalized on the untreated samples, obtained by END-seq (D) and S1-END-seq (E) in untreated and dye + light treated cells recovered for the indicated times. U2OS cells used as a control of cell line harboring elevated telomeric SSBs and DSBs.

**Supplementary Figure 3.**
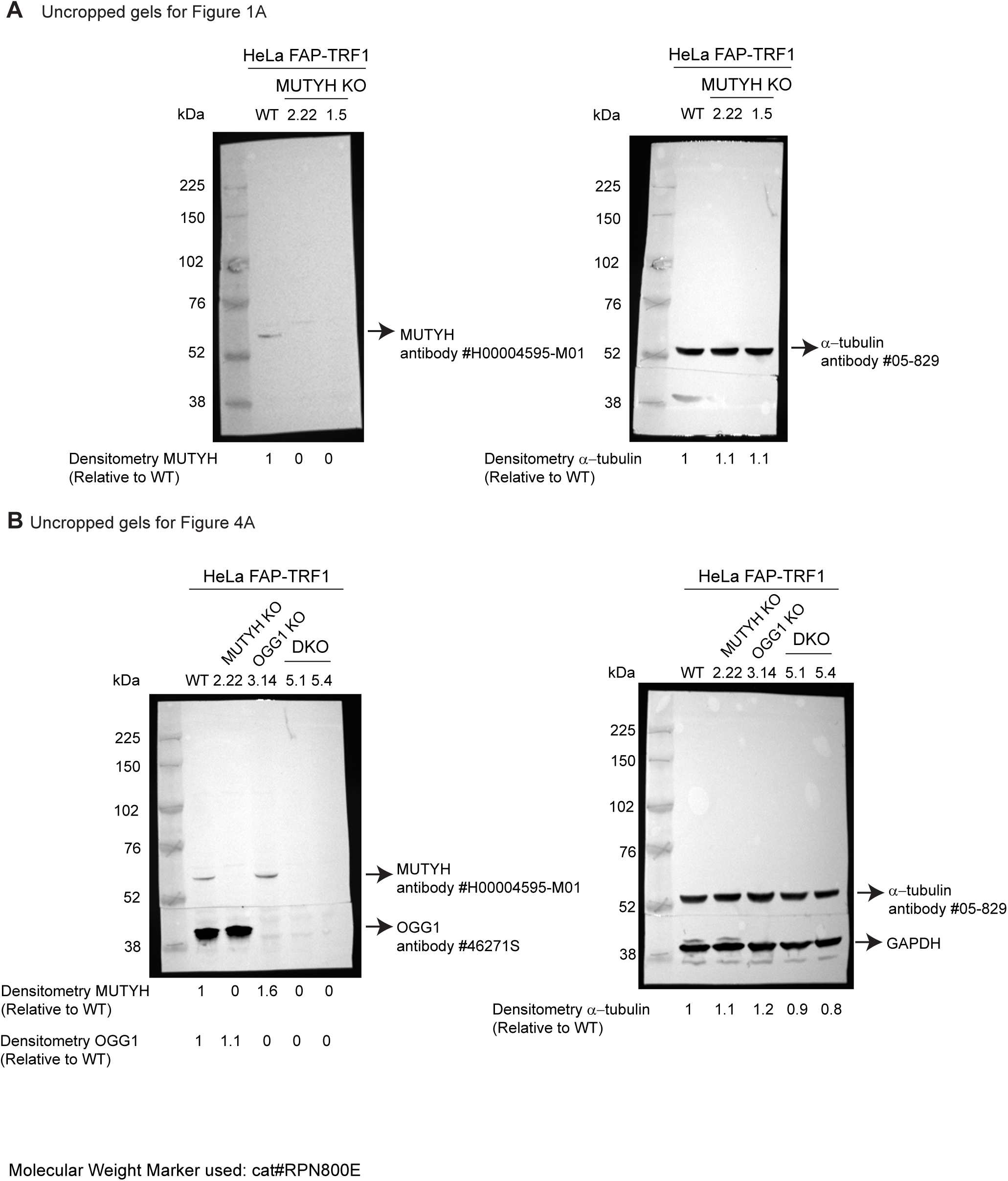

